# Global DNA signatures of temperature and nutrient limitation in prokaryotes

**DOI:** 10.1101/2025.01.13.632720

**Authors:** Tomer Antman, Ohad Lewin-Epstein, Tamir Yerushalmi, Yonatan Broder, David Zeevi

## Abstract

Microorganisms adapt to their environment through changes in both their genes and overall genome composition. However, identifying universal principles of genomic adaptation remains challenging because different environments present multiple, overlapping selective pressures. To overcome this challenge, we analyzed DNA composition patterns across diverse environments using machine learning. By examining tetranucleotide frequencies from 1,112 marine and soil metagenomic samples, we discovered that environmental temperature can be accurately predicted from DNA composition alone (R^2^=0.82). This temperature signal remained robust even when analyzing individual bacterial phyla and classes, suggesting a fundamental adaptive response. To examine this adaptation mechanism, we analyzed GC content relationships with temperature and found opposing relationships in different environments: positive correlations in soil but negative correlations in marine samples. We show that this inverse relationship in marine environments is driven by nutrient availability, as GC content increases with nutrient levels while nutrients decrease with temperature in marine ecosystems. Despite these contrasting GC patterns, we identified specific tetranucleotides composed of equal numbers of GC and AT bases (50% GC content) that showed consistent temperature correlations across all environments. These findings reveal a complex interplay between temperature adaptation and nutrient limitation in shaping microbial genomes.

## Introduction

A key challenge in microbial ecology is to unravel the complex interplay between environmental pressures and their effects on microbial diversity, interactions, and evolution^1–4^. The genomic effects of these pressures can be broadly categorized into two types. The first is selection of functions that increase organismal fitness, operating through adaptive genetic changes in protein-coding genes or regulatory regions. For instance, antibiotic-resistance genes confer survival advantages in the presence of antibiotics^5^, while cold-shock proteins are favored in environments with temperature fluctuations^6^. The second type of effect operates on the scale of the entire genome, influencing the composition of nucleic and amino acids. Examples of this genome-wide effect include the prevalence of GC-rich genomes in hyperthermophiles, which helps maintain DNA stability at high temperatures^7^, or genome streamlining in nutrient-poor environments as an evolutionary pressure for minimizing resource utilization^8^.

Understanding how genome-wide adaptations to environmental pressures emerge across diverse environmental contexts remains a significant challenge. Two main approaches exist for studying these adaptations: conducting laboratory evolution experiments and analyzing sequences from natural environments. Laboratory evolution experiments, while controlled, are limited because genome-wide selective pressures are often too weak to manifest within experimental timescales. Environmental sequence analysis is challenging due to the complexity of obtaining and analyzing data from diverse ecosystems, as well as disentangling the effects of multiple selective pressures acting simultaneously.

Global shotgun metagenomic surveys offer a unique opportunity to study the impact of environmental selective pressures on microbial genomes^9–12^. These studies simultaneously collect DNA sequences and associated environmental parameters, including temperature, pH, and the availability of nutrients such as nitrogen, which are dominant selective pressures acting on microbial communities^9,13–16^. Environmental sampling inherently captures the microbial communities most adapted to local conditions, revealing key genomic features that respond to these selective pressures. However, most studies to date have focused on describing these pressures within a single environmental context, such as marine or soil ecosystems^9,14,17,18^, due to two main challenges. First, different environments harbor distinct microbial communities with varying genomic compositions—with less than 1% of bacterial species considered common and prevalent across biomes^19^—making inter-environmental comparisons difficult. Second, the methods and units used to measure environmental parameters often differ across ecosystems. For instance, nitrate concentrations in marine environments are typically measured in μmol/L^20^, while in soil they are often reported in mg/kg^21^. One promising approach to overcome these obstacles is to focus the analysis directly at the DNA sequence level, bypassing the need for genome- or gene-centric comparisons. Additionally, selecting environmental parameters that can be consistently measured across diverse ecosystems, such as temperature, can facilitate more robust cross-environment analyses.

In this study, we characterize how fundamental environmental conditions and stressors, such as temperature and nutrient limitation, shape the DNA composition of microbes across diverse ecosystems. To detect these potentially subtle and complex signatures, we employed regression-based machine learning algorithms to analyze tetranucleotide frequencies (4-mers) from 1,112 metagenomic samples across soil and ocean microbiomes^10–12,17,22,23^. Our machine learning model successfully inferred temperature with high accuracy, greatly surpassing the predictive power of GC content, which correlated positively with temperature in soil samples but negatively in marine samples. Since 4-mer frequencies are known to be useful in determining taxonomy, we wanted to examine whether the signal might be primarily driven by phylogeny. We demonstrated that temperature inference remained robust even when restricted to sequences from a single phylum or class. To understand whether these 4-mer patterns reflect selection at the DNA level rather than protein-coding constraints, we examined synonymous codon usage. The abundance of high-GC codons followed the same pattern as overall GC content: positive correlation with temperature in soil and negative in marine environments. We propose that this discrepancy in marine samples is due to nutrient limitation, particularly nitrogen and phosphorus, which are negatively correlated with temperature in marine ecosystems^24,25^. Supporting this hypothesis, marine samples with low variance in nutrient content exhibited a positive correlation between high-GC codon usage and temperature, resembling the pattern observed in soil. These findings highlight the interplay between two genome-wide selective forces: temperature adaptation and nutrient limitation, with nutrient availability exerting a stronger influence on genomic composition in marine environments. Our findings uncover subtle yet pervasive signatures of adaptation in metagenomic data that significantly deepen our understanding of how environmental pressures shape microbial genomes across diverse ecosystems.

## Results

### Temperature is accurately inferred from metagenomes across environments

To examine the effect of abiotic parameters on microbial communities across environments, we compiled 1,112 unique marine and soil metagenomic samples (**Fig. 1a**). This collection includes 490 soil metagenomic samples from the National Ecological Observatory Network^11,26^ (NEON) and 622 marine metagenomic samples from three datasets: bioGEOTRACES^10^ (n = 472), the Tara Oceans consortium^17^ (n = 128), and the POLARSTERN cruise^12^ (n = 22; used for model validation). These datasets span a wide array of oceanic and land biomes and diverse environmental conditions (**Fig. 1a**, **S1** and **S2**). For each metagenomic sample, we also obtained its corresponding metadata, including the sampling time and location, temperature, pH, and other environmental parameters (**Table S1**).

**Figure 1:**
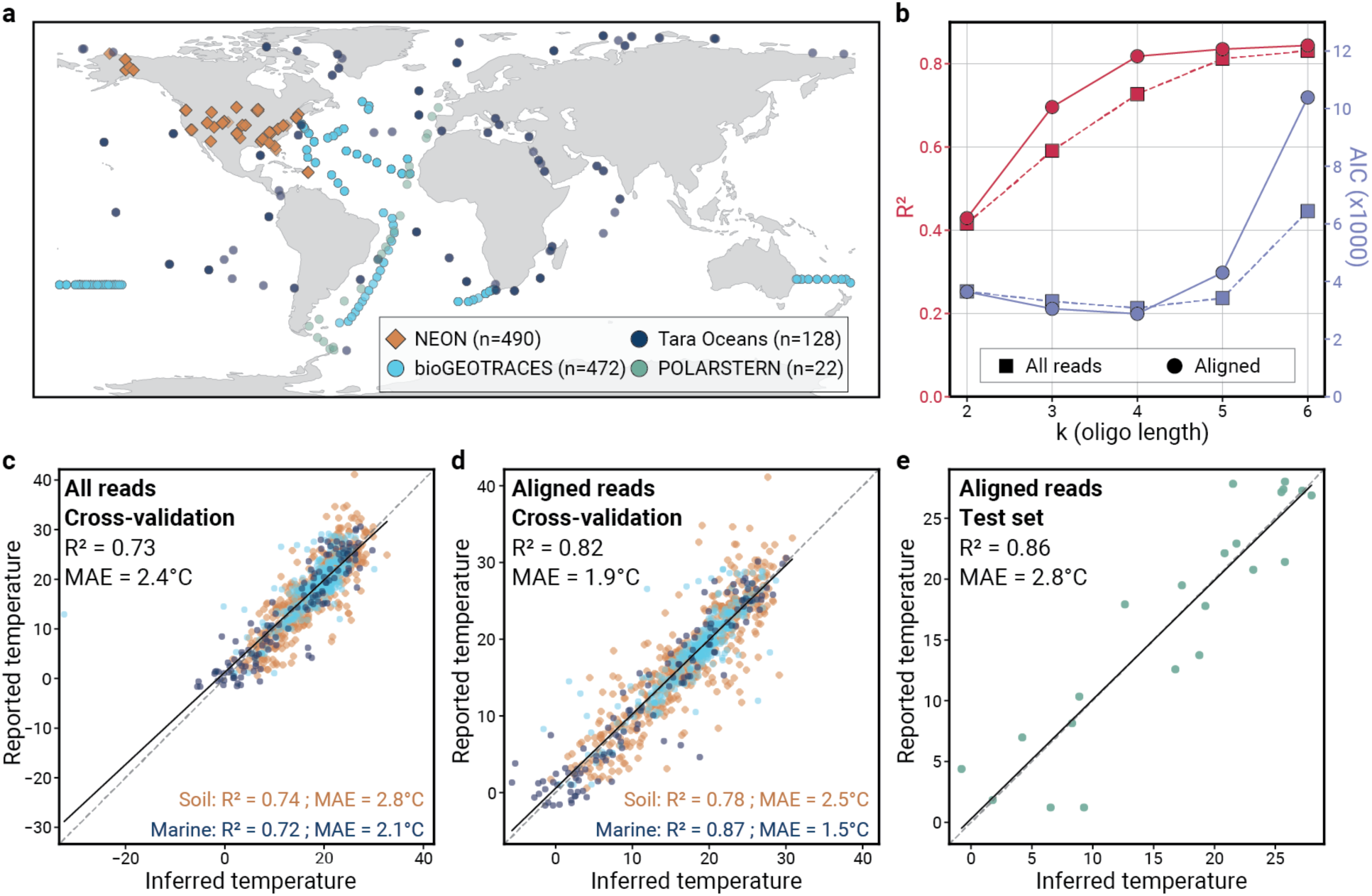
Temperature is accurately inferred from global metagenomes of diverse origin. **[a]** A world map of sampling locations in the curated datasets. **[b]** Model quality assessment of k-mer size (x-axis) based on R² (left y-axis; red) and Akaike Information Criterion (AIC; right y-axis; blue). Higher R² and lower AIC are typical of better models. The plot compares unaligned reads (squares, dashed lines) and reads aligned to reference genes (circles, solid lines). **[c-e]** Scatter plots showing reported temperatures (y-axis) versus temperatures inferred from 4-mer frequencies (x-axis), with colors matching the legend in panel [a] (solid line, linear regression; dashed line, identity). **[c]** Leave one site out cross-validation (LOSO-CV) using 1M reads per sample. **[d]** LOSO-CV using 1M reads aligned to prokaryotic gene catalogs per sample. **[e]** Inference on an independent test set using 1M aligned reads.

We used this broad dataset to investigate how DNA compositional signatures vary across temperature gradients in diverse environments. Temperature at sampling time was the only parameter comprehensively measured across these soil and marine samples, spanning a range of −1.6 - 30.6°C in marine samples and 0.6 - 44.0°C in soil (**Fig. S1** and **S2**) Furthermore, temperature is a fundamental abiotic factor that regulates numerous biological processes, including DNA replication and cell growth^9,15^.

To infer temperature from the DNA composition of microbial communities we focused on DNA k-mer frequencies as features for inference, as they can efficiently capture broad DNA composition patterns, they are present across environments, and comprise a small number of features compared to the number of genes or species. We used a Bayesian Ridge regression model—a simple regularized linear model—to improve its interpretability and reduce the potential of overfitting.. Our model was trained using the NEON, bioGEOTRACES, and Tara Oceans datasets (n = 1090) and we applied leave-one-site-out cross-validation (LOSO-CV, Methods) to ensure that it is not confounded by geographical proximity or sampling bias. In our analysis, 4-mers were shown to be superior to other oligo lengths in terms of balancing between goodness of inference fit (R^2^) and relative model quality (Akaike Information Criterion, AIC^27^; **Fig. 1b**).

First, we predicted sample-associated temperatures using the 4-mer DNA composition of raw metagenomic samples. To control for varying sequencing depths, the 4-mer vectors of each sample (136 features, considering reverse complement 4-mers) were constructed using only one million reads. This LOSO model predicted the temperatures measured at sampling with an R^2^ of 0.73 and a mean absolute error (MAE) of 2.4°C (**Fig. 1c**).

Hypothesizing that we could improve these results by focusing on reads from coding regions, which are typically under stronger environmental selection, we conducted a prokaryotic gene-focused analysis by aligning the reads to prokaryotic gene catalogs^4,9^ (Methods) and analyzing one million mapped reads for each sample. In these analyses, we considered only the sense strand 4-mers (256 features, without considering reverse complements). The prokaryotic gene-focused model showed an even greater temperature inference accuracy, reaching an R^2^ of 0.82 and MAE of 1.9°C (**Fig. 1d**). This improvement supported our hypothesis regarding selection in coding regions, and we therefore used this gene-focused approach in subsequent analyses.

Given the differences in heat conductance and temperature fluctuations between soil and water, we sought to determine the differences in inference quality between environments. When analyzing the performance of the model on ocean and soil samples separately (training on ocean and soil samples combined, but evaluating on ocean and soil separately), we obtained an R^2^ of 0.87 and MAE of 1.5°C for ocean samples, and R² of 0.77 and MAE of 2.5°C for soil samples (**Fig. 1d**). Surprisingly, these results are similar to the performance of models trained and evaluated separately on ocean and soil samples (R^2^ = 0.88, MAE = 1.6°C and R^2^ = 0.79, MAE = 2.5°C respectively; **Fig. S3**). Since microbial communities adapt to environmental changes at varying rates that may lag behind temperature fluctuations, we hypothesize that our model performs better in marine environments which are more thermally stable and likely contain less relic DNA^28^.

To assess the robustness of the model to external datasets collected and sequenced by different labs, we tested its performance by inferring the temperatures of samples from the POLARSTERN cruise. The POLARSTERN samples span across wide ranges of both latitudinal coordinates (between 62°S and 47°N) and measured temperatures (between 1.2°C and 28.0°C), all from the same depth (20m)^12^. The model predicted the samples’ temperatures with an R^2^ of 0.86 and MAE of 2.8°C (**Fig. 1e**). These results demonstrate that temperature at sampling time can be robustly inferred from microbial DNA composition alone, suggesting a strong impact of the temperature on the DNA features of the resident microbial community.

Having found a stronger temperature signal in coding regions, we sought to determine whether this signal could be attributed to codon usage or amino acid composition. To this end, we built new models based on codon and amino acid frequencies, calculated from gene-aligned reads. These models achieved lower accuracies than gene-aligned 4-mer models (R^2^ = 0.75, MAE = 2.4°C for codons and R^2^ = 0.63, MAE = 3.2°C for amino acids; **Fig. S3**). The superiority of the 4-mer model could stem from including more parameters, and specifically capturing some information on neighboring codons. Nevertheless, these results suggest that the temperature signal caught by the microbiome extends beyond the composition of amino acids or the codon usage.

To put the temperature inference ability of the model in context, we examined the accuracy that could be achieved from an alternative analysis, based on geographic parameters, including latitude, depth, and temperature at the sea surface (collected from NASA^29^; Methods). This alternative inference achieved R^2^ = 0.91 and MAE = 1.3°C on marine samples (**Fig. S4**) while our 4-mer model applied on marine samples alone obtained slightly lower yet comparable results of R^2^ = 0.87 and MAE = 1.5°C (**Fig. 1d**). This suggests that the accuracy achieved by 4-mers approaches the limit set by the most informed model that we managed to create based on remote sensing data. Altogether, these results demonstrate that the environmental temperature can be robustly inferred from microbial DNA composition alone, suggesting a strong impact of the temperature on the DNA features of the resident microbial community.

### Phylum- and class-level phylogeny does not confound temperature inference

Tetranucleotide (4-mer) frequencies have long served as a powerful tool for taxonomic classification, traditionally used to distinguish between microbial taxa based on their distinct genomic signatures^30,31^. Typically, these frequencies are leveraged to delineate boundaries between microbial groups, with each taxonomic lineage characterized by its unique 4-mer composition. Given this established paradigm of using 4-mer frequencies as a phylogenetic discriminator, we sought to investigate whether our observed temperature-associated genomic patterns might reflect the underlying taxonomic structure rather than a broader environmental adaptation mechanism. We thus systematically examined the temperature inference capabilities within individual phyla and classes to validate our temperature inference model separately from these taxonomic classifications.

We used the GMGC^4^ and OM-RGC^9^ datasets to assign each read with its taxonomy. Due to the limited number of metagenomic reads per taxon, we predicted temperature using 4-mers from 5,000 reads for each taxon in each sample (Methods). In this analysis, we trained temperature prediction models, for each taxon that had 5,000 reads in at least 30 samples per dataset (Methods; example for phylum Actinomycetota with R² = 0.72 and MAE = 2.7°C; **Fig. 2a**; yellow stars in **Fig. 2b, c**; **Fig. S5**). We hypothesized that if the temperature predictions were merely reflecting taxonomic composition—since different taxa might be adapted to different temperatures—then focusing on a single taxon at a time should weaken the predictive signal. For comparison, we created a null distribution by sampling 5,000 random reads from each sample, using the same number of soil and marine samples in which each taxon was present, and repeated this process 1,000 times. We compared each phylum and class to this background distribution (**Fig. 2b, c**; **Fig. S5**). In contrast to the null hypothesis that temperature predictions reflect taxonomic compositions, the R² values of the taxon-specific inferences (yellow stars) are typically higher than the null distribution. This improved performance may reflect reduced noise when analyzing phylogenetically coherent reads compared to mixed-taxon sequences. By demonstrating temperature prediction within each phylum and class independently, we showed that the observed signal is not driven by a single dominant phylum or a small set of taxa. Instead, the temperature-associated DNA composition signal is detectable across diverse phylogenetic groups, regardless of their relative abundance in the samples.

**Figure 2:**
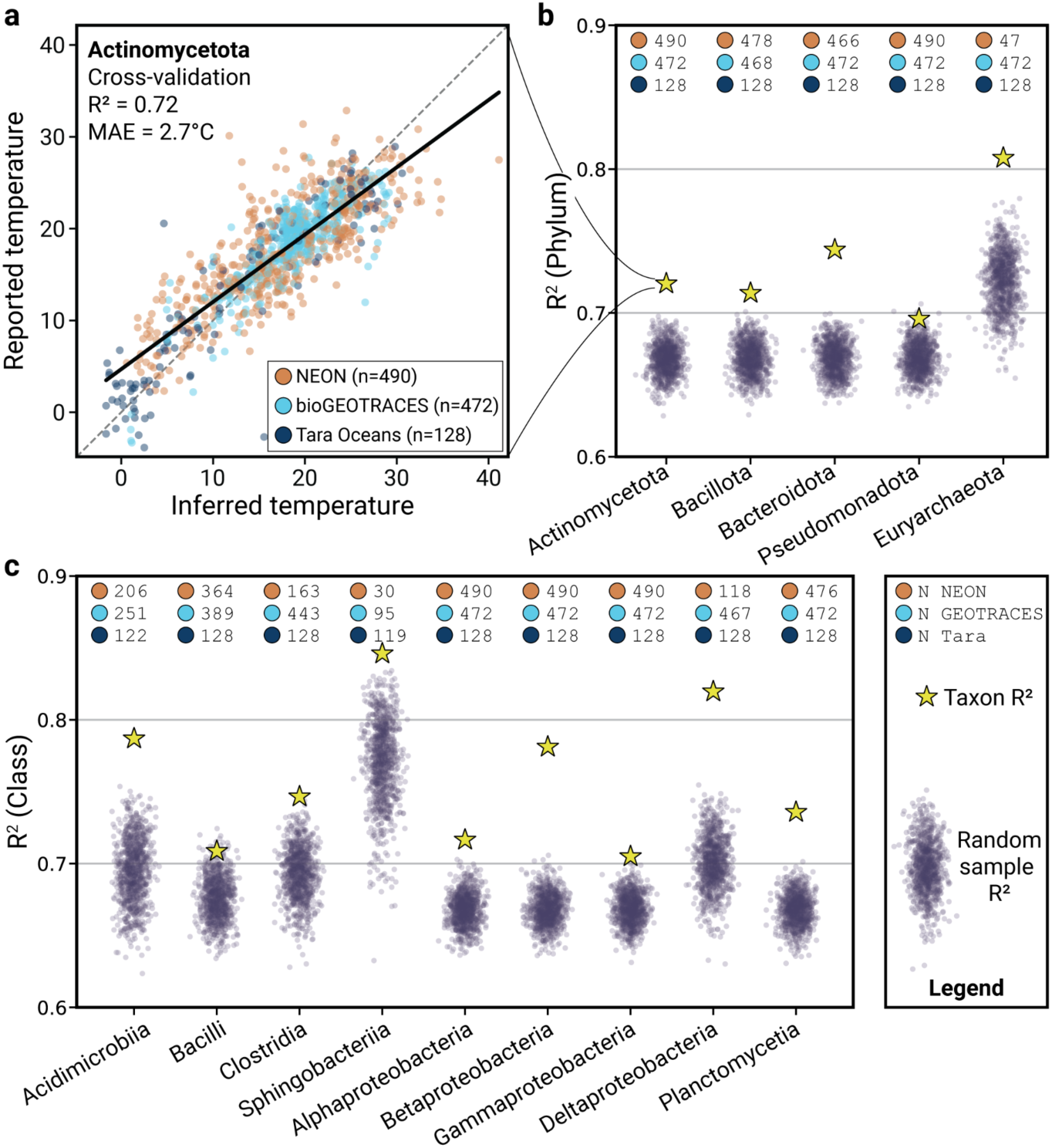
Temperature inference remains robust within individual phyla and classes. **[a]** Scatter plot of inferred versus reported temperatures using cross-validation and 4-mers from metagenomic reads assigned to Actinomycetota. **[b, c]** Jitter plots depict the R² of temperature inference from sample- and read-randomized sampling. Yellow stars represent the R² for temperature inference using 5,000 reads mapped to each taxon within different phyla **[b]** and classes **[c]** per sample. Per-dataset sample counts are shown above each box plot.

We note that reads assigned to some taxa exhibit higher temperature inference abilities than others. In some cases, such as with Euryarchaeota (**Fig. 2b** and **S5e**) or Sphingobacteriia (**Fig. 2c** and **S5i**) this is likely due to the exclusion of soil samples in which they are not present and where there is an overall lower inference quality (**Fig. 1d**, **Fig S3**). However, Betaproteobacteria-assigned reads have higher inference goodness of fit than Alpha- and Gammaproteobacteria, despite being present in all samples (**Fig. 3c**, **S5j-l**). Overall, this analysis demonstrates that temperature-associated 4-mers represent a pervasive ecological adaptation signature, transcending taxonomic boundaries. This underscores the fundamental role of the environment in shaping microbial genomic composition.

**Figure 3:**
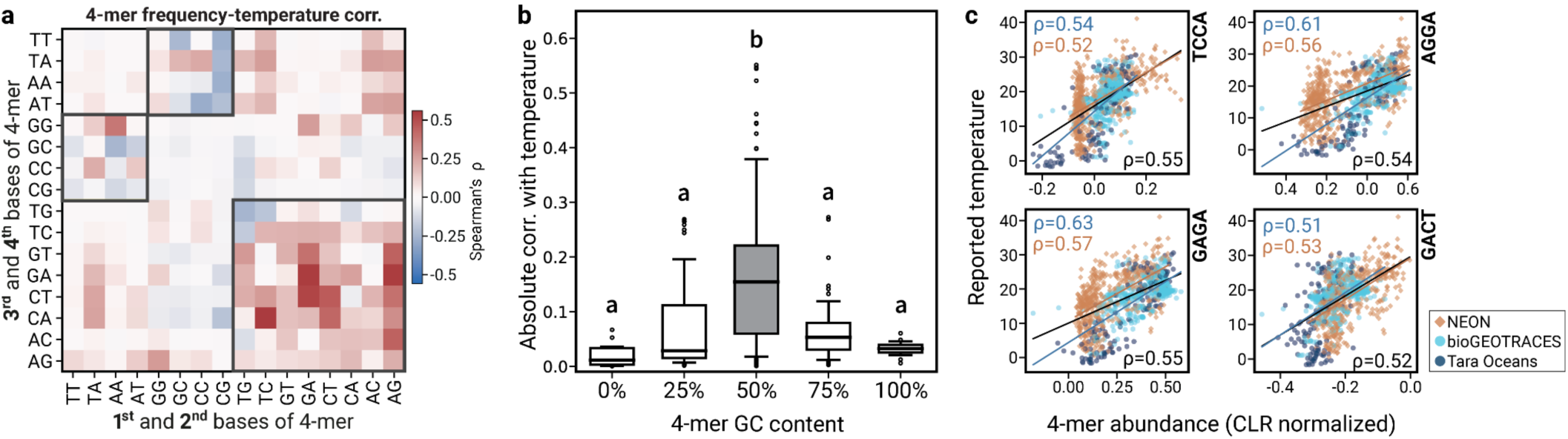
Temperature is consistently correlated with 50% GC content 4-mers in marine and soil biomes. **[a]** Spearman correlations between temperature and CLR-transformed 4-mer frequencies. 50% GC 4-mers highlighted by gray squares. **[b]** Box plots (line, median; box, interquartile range; whiskers, 10^th^ and 90^th^ percentiles) of the absolute correlation values between temperature and CLR-transformed 4-mer frequencies, grouped by GC content of the 4-mers. Different letters above the boxes indicate statistically significant differences between groups (Tukey’s HSD test, p < 0.05). Groups sharing a letter are not significantly different from each other. **[c]** The 4 CLR-transformed 4-mer frequencies that were best correlated with temperature. Overall correlation (black), soil correlation (orange), and marine correlation (blue).

### Temperature significantly correlates with 50% GC 4-mers across biomes

To better understand how our 4-mer model achieved robust predictions (**Fig. 1d**), we analyzed the relationship between individual 4-mer frequencies and temperature. We hypothesized that some 4-mer frequencies were more robust than others to the difference between the environments. We therefore examined the correlation of each center log ratio (CLR)-transformed 4-mer frequency with temperature across marine and soil biomes. We observed that 4-mers with 50% GC content (exactly two G/C bases) exhibited significantly higher positive or negative correlations with temperature than non-50% GC content 4-mers (Mann-Whitney *U* p < 10⁻¹⁰; **Fig. 3a, b**).

Among these features, we found that the frequencies of several specific 4-mers showed particularly strong correlations with temperature across environments. For example, AGGA exhibited a high correlation with temperature in all samples combined (Spearman’s ρ = 0.54, p < 10⁻¹⁰; **Fig. 3c**). The correlation was strong in both marine (ρ = 0.61) and soil (ρ = 0.56) environments. Similarly, TCCA, GAGA, and GACT showed consistent correlations across biomes (**Fig. 3c**). This suggests that the frequencies of 50% GC 4-mers may be genomic markers for temperature that are robust across environments.

### Codon GC content associates with temperature and confounded by nutrient availability

Given that 4-mers with 50% GC content showed the strongest correlations with temperature, we next investigated the broader relationship between GC content and temperature in our samples. While GC content has been previously associated with temperature, particularly in thermophiles where it is a well-established method of thermal adaptation^7,32,33^, our findings about 50% GC 4-mers suggested a more nuanced relationship might exist. As a first step, we correlated the GC content of metagenomic samples (using gene sequence-aligned DNA reads) with temperature at the time of sampling. We found a negative correlation in marine samples (Spearman’s ρ = −0.42, p < 10^−10^; **Fig. 4a**), and a positive correlation in soil samples (Spearman’s ρ = 0.55, p < 10^−10^; **Fig. 4a**). These opposing relationships between GC content and temperature in different environments explain why 4-mers with balanced GC content emerged as the most robust temperature predictors (**Fig. 3a, b**).

**Figure 4:**
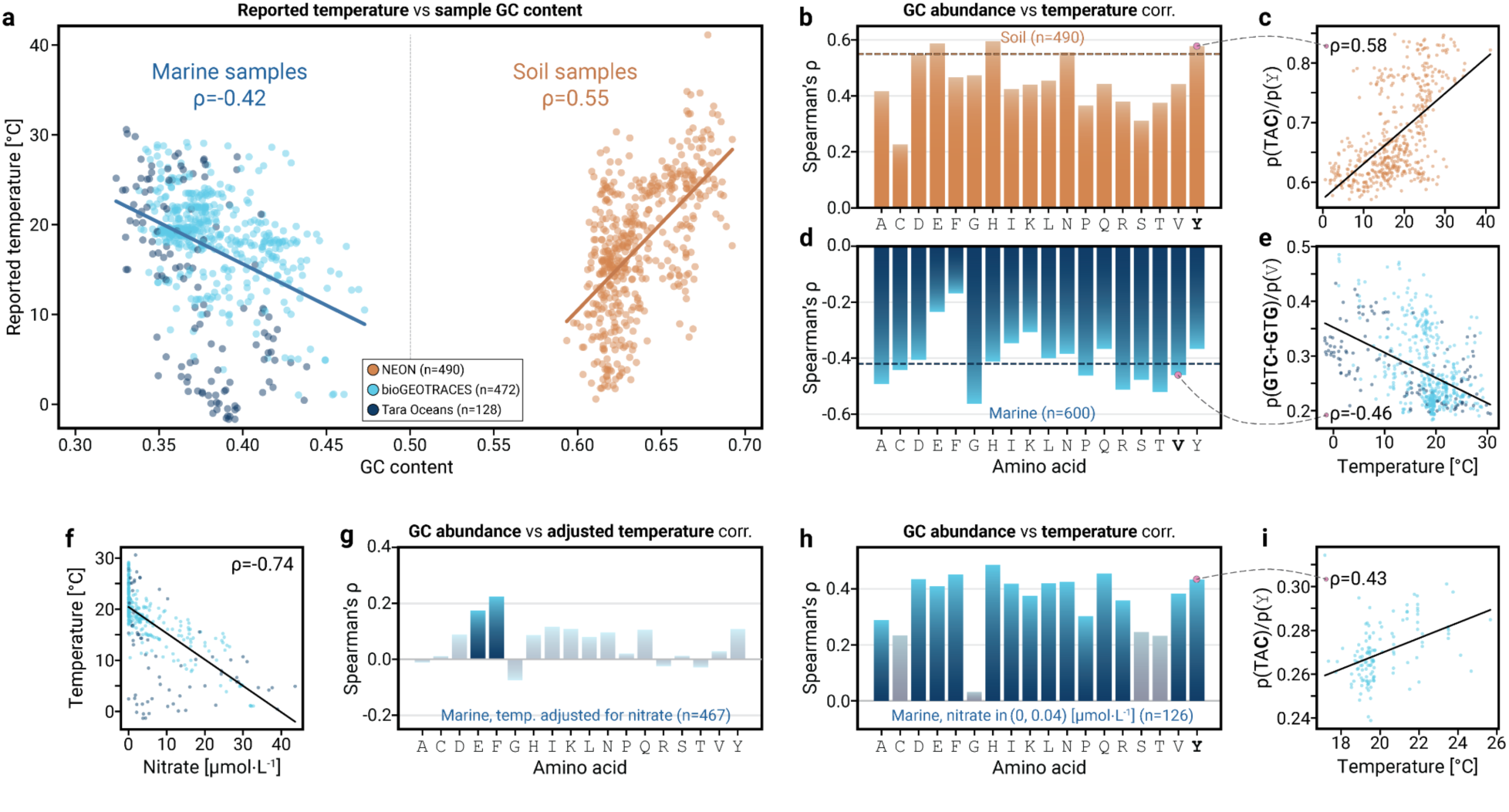
GC-rich codons present conflicting temperature trends in soil and water. **[a]** Temperature plotted against GC content for marine (blue) and soil (orange) samples. ρ, Spearman’s correlation coefficient. **[b]** Spearman correlations between temperature and amino acid GC-fraction for soil samples. The overall temperature-GC correlation in this environment is shown as a colored dashed line. **[c]** an example of the correlation between temperature and amino acid GC-fraction, for tyrosine in soil samples. **[d]** Same as [b], for marine samples. **[e]** Same as [c] for valine in marine samples. **[f]** Temperature plotted against nitrate concentrations in marine samples. **[g]** Same as [b], for marine amino acid GC-fraction-temperature correlations after adjusting for nitrate effects. Non-significant correlations are faded (Bonferroni corrected p-value < 0.05). **[h]** Same as [b] for marine amino acid GC-fraction-temperature correlations in samples with nitrate in the range of 0-0.04 μmol/L (non-inclusive; n = 126). **[i]** Same as [c] for tyrosine in nitrate-limited samples.

To understand whether these 4-mer patterns reflect selection at the DNA level rather than protein-coding constraints, and to further investigate the contrasting GC-temperature relationships we observed between environments, we examined synonymous codon usage. To this end, we calculated, for each amino acid, the relative frequency of its codons with the highest GC-content. For example, in tyrosine, we calculated, in each sample, the frequency of the TA<u>C</u> codon, normalized to the frequencies of TA<u>C</u>+TA<u>T</u>. We found that this relative frequency of GC-abundant codons (GC codon bias) is correlated positively with temperature in soil samples (**Fig. 4b**; example for tyrosine in **Fig. 4c**), and negatively in marine samples (**Fig. 4d**; example for valine in **Fig. 4e**). In some amino acids, the correlation of temperature with the amino acid GC-fraction exceeded the correlation of temperature with overall metagenomic GC-content (**Fig. 4b, d**, dotted line).

GC content has been shown to increase with ocean depth along with inorganic nitrogen^34,35^, while water temperature exhibits the opposite behavior^36^. This negative correlation between temperature and nutrient availability, caused by the photic-gradient structure of the marine environment, was also observed in our analysis (**Fig. 4f** for nitrate; **Fig. S6** for phosphate and depth). The strong correlation between temperature and nitrate (Spearman’s ρ = −0.74) might suggest that the GC codon bias in marine environments (**Fig. 4a, c**) is also constrained by nutrient availability. To test this hypothesis, we performed a two-step residual analysis by fitting a linear regression of temperature against nitrate or phosphate and calculating the residuals (Methods).

We found that correlations between temperature residuals and GC codon bias become mostly insignificant (faded bars) with some correlations changing signs (**Fig. 4g, S6** for nitrate; **Fig. S6** for phosphate; **Table S2**). Additionally, we performed an analysis of covariance (ANCOVA) to assess the relationship between the GC codon bias and temperature while controlling for nutrient availability and arrived at similar results (**Table S3**). This suggests that the negative correlation between temperature and codon GC-content in marine environments was likely driven by nutrient availability.

We hypothesized that the intrinsic relationship between GC codon bias and temperature might be positive in marine environments as well, but that this relationship is masked by the strong negative correlation between nutrients and temperature, reminiscent of Simpson’s paradox^37^—a statistical phenomenon where a trend present in different groups disappears or reverses when these groups are combined. To test this hypothesis, we recalculated the Spearman correlations between temperature and the GC codon bias using only samples with nitrate and phosphate concentration in ranges that were uncorrelated with temperature (**Fig. 4h, S6** for nitrate; **Fig. S6** for phosphate). In these subsets, where nitrate’s confounding effect is naturally controlled for, the correlation pattern either becomes non-significant or inverts compared to the original correlation between temperature and GC codon bias (**Fig. 4d**). This inverted association resembles the trend observed in soil samples (**Fig. 4b**). However, the overall GC content correlation in these ranges remained negative or became insignificant. This insight is supported by previous studies showing that GC content at third codon positions is more likely to be correlated to temperature than genomic GC in prokaryotes^38^. Our finding supports the hypothesis that in marine environments, nutrient availability is the primary driver of codon GC content variation, while temperature acts as a secondary influence, suggesting these environmental factors exert compound selective pressures on coding-sequence GC composition.

## Discussion

In this study, we analyzed DNA-composition signatures in metagenomic samples from soil and marine environments. Our analysis revealed a universal signal of temperature adaptation that transcends ecosystem boundaries. This temperature signature persists even when controlling for phylum and class-level taxonomy, providing compelling evidence for a fundamental response that is apparent at the DNA sequence level. Furthermore, we uncovered a complex interplay between temperature and nutrient availability in shaping codon usage, particularly in marine ecosystems. These findings significantly deepen our understanding of how global-scale environmental pressures shape microbial genomes across diverse habitats, highlighting the potential of leveraging large-scale metagenomic datasets to uncover subtle yet pervasive signals of environmental adaptation.

It is important to acknowledge the inherent complexity in interpreting tetranucleotide frequencies, which likely encompass multiple layers of biological information, including residual phylogenetic signatures, codon usage patterns, and DNA and RNA stability factors. Despite our findings on phylogeny and codon usage, the ability to infer temperature from tetranucleotides likely reflects the cumulative influence of multiple evolutionary pressures and molecular mechanisms. This multifaceted nature presents challenges in fully isolating individual contributors to temperature adaptation, and their relative importance may vary across environments or taxa. Future studies analyzing the contribution of specific genomic features such as RNA secondary structures to temperature adaptation could potentially provide additional insights into microbial temperature adaptation at the genomic level^38–40^.

The strong correlation between 4-mer frequencies and temperature raises questions about causality and adaptation mechanisms. While multiple interpretations are possible, one plausible explanation is that 4-mer signatures reflect ecological dynamics where certain bacterial strains perform better than others under specific temperature conditions. These successful strains may carry genomic adaptations that have evolved over time in response to their thermal environment, potentially explaining the consistent DNA compositional patterns we detected. However, we cannot rule out alternative explanations, such as indirect effects of temperature on community composition through other environmental factors, or the possibility that some 4-mer patterns arise from neutral processes that happen to correlate with temperature.

The observation of what appears to be Simpson’s paradox in the relationship between amino acid GC content and temperature suggests a complex interplay between nutrient availability and thermal adaptation. When controlling for nutrient concentrations, particularly in marine environments, the codon GC-temperature relationship shifts dramatically, potentially indicating that nutrient availability may constrain the evolutionary trajectories available to microorganisms. While several interpretations are possible, one hypothesis is that although temperature creates selective pressure for specific genomic features, the ability of microorganisms to evolve these adaptations could be fundamentally limited by nutrient availability. For instance, the energetic and material costs of maintaining GC-rich genomes—which require more nitrogen than AT-rich alternatives— might restrict the evolution of temperature-optimal genomic compositions in nutrient-poor environments. If our interpretation is correct, it would suggest that the evolutionary “solution space” for adaptation may be narrowed by resource limitations, potentially forcing microorganisms to adopt alternative genomic strategies. These findings highlight the importance of considering multiple environmental factors when studying adaptation, as the response to one selective pressure may be shaped by constraints imposed by others.

Importantly, our analysis captures community-level genomic signatures that may not directly translate to individual bacterial genomes or their optimal growth temperatures. Any given strain within these communities might thrive at temperatures different from those where it was found, but their presence indicates superior competitive fitness relative to other microbes under the actual environmental conditions. This distinction between optimal competitive temperature and physiological temperature optimum highlights a crucial aspect of microbial ecology—the difference between what is optimal for an organism in isolation versus what it can successfully endure in community dynamics with competition in nature. Our findings thus underscore the importance of in situ metagenomic analyses for understanding real-world microbial adaptation, as laboratory-based temperature optima studies may not fully capture the competitive dynamics that shape microbial communities in their natural habitats.

The potential impact of temperature adaptation on global nutrient cycles warrants consideration. A simple calculation illustrates this point: if microbial GC content were to increase by 0.5% per degree Celsius (as observed in soil environments), each GC pair requiring one additional nitrogen atom compared to an AT pair, a typical microbe with a 2 million base pair genome would require 10,000 additional nitrogen atoms per degree of warming. Given an estimated 10³⁰ microbial cells on Earth^41^, this would translate to approximately 10³⁴ additional nitrogen atoms, or roughly 250,000 tons of additional nitrogen sequestered in microbial DNA alone per degree Celsius. While this quantity may seem modest compared to the global nitrogen cycle, which processes hundreds of millions of tons annually, it highlights how genomic adaptation to temperature could influence nutrient dynamics at a global scale. These calculations become particularly relevant in the context of anthropogenic climate change and increasing nitrogen deposition. Our findings suggest that the microbial response to warming may vary significantly between environments based on nutrient availability, potentially creating feedback loops between temperature adaptation, nutrient cycling, and microbial community composition. Understanding these relationships is crucial for predicting ecosystem responses to climate change, as shifts in microbial genomic composition could affect both their functional capabilities and their nutrient requirements. However, given the complex relationships we observed between temperature, nutrients, and genomic composition, predicting these responses will require careful consideration of local environmental conditions and constraints.

## Methods

### Datasets curation

#### Samples

Marine microbiome samples were downloaded from ENA with accessions ENA:PRJEB1787 (Tara Oceans prokaryotic fraction), ENA:PRJEB9740 (Tara Oceans Polar Circle prokaryotic fraction), ENA:PRJNA385854 (bioGEOTRACES), and ENA:PRJEB34453 (POLARSTERN). Soil samples were downloaded from the National Ecological Observatory Network (NEON) website through the links provided in data product ID:DP1.10107.001 (Soil microbe metagenome sequences)^11^.

#### Environmental variables

Tara Oceans metadata was downloaded from PANGAEA with accession number PANGAEA.875576^42^. To improve our confidence in nitrate measurements, we examined sensor data submitted to ENA and discarded 26 samples with irregular nitrate sensor reports (< 0 [µmol/L]). bioGEOTRACES metadata was compiled from the GEOTRACES intermediate data product v.2^43^. NEON metadata that appears in data product ID:DP1.10086.001 (Soil physical and chemical properties, periodic) was downloaded from the NEON website^26^. As bioGEOTRACES and Tara Oceans reported nitrate measurements in different units (µmol/kg and µmol/L, respectively), we standardized the values by converting the bioGEOTRACES nitrate values to µmol/L by multiplying them by the density of seawater (1.021 kg/L)^44^.

### Quality trimming

Data quality control and filtering/trimming tools include Trimmomatic^45^ and fastp^46^. We applied Trimmomatic 0.36 with the following command line parameters: [*PE*, *phred33*, *LEADING:* 10, *TRAILING:* 10, *SLIDINGWINDOW:* 4:15, *MINLEN:* 50]. We applied fastp with the following command line parameters: [*length_required:* 100, complexity_threshold: 20, *dedup*, *trim_poly_g*, *detect_adapter_for_pe*, *low_complexity_filter*, *overrepresentation_analysis*]. We rarified all samples to 1 million reads and filtered out samples with fewer reads.

### Gene-catalog alignment

Using bowtie2^47^, marine and soil samples were aligned to the Ocean Microbiome Reference Gene Catalog (OM-RGC)^9^ and the Global Microbial Gene Catalog (GMGC)^4^, respectively. Reads with no gene mapping were filtered out of the data. In analyses using aligned reads, we rarified all samples to 1 million aligned reads and filtered out samples with fewer reads.

### Tetranucleotide frequency calculation

For every read, we counted the occurrences of unique tetranucleotides. The counts of the paired-end files were summed and converted to frequencies, thus creating per-sample tetranucleotide frequencies (4-mers). For reads aligned to reference gene sequences, where it was important to consider the specific DNA strand from which the read originated, a 256-feature vector was created. For general, unaligned reads we considered the reverse complemented 4-mers (e.g., “AAAA” and “TTTT” were considered as “AAAA/TTTT”). This process resulted in a 136-feature vector. Lastly, to address the compositional nature of the data, we applied centered log-ratio (CLR) transformation to the 4-mer frequency vectors.

### Regression model and cross-validation

Bayesian Ridge regression was performed using the default “Scikit-learn”^48^ settings. The model was assessed using a form of “Leave-One-Group-Out” (LOGO) cross-validation. To avoid locality-driven over-training, the marine samples were grouped by geolocation, and the soil samples were grouped by field site. This way, the inference was ignorant of the sample’s source, depth, and surroundings. We named this stratification “Leave-One-Site-Out” (LOSO). Sample groupings are presented in **Table S1**. The difference in the Akaike Information Criterion (AIC) between the models was calculated as 𝛥𝐴𝐼𝐶 = 2𝑘 + 𝑛 ⋅ 𝑙𝑛(𝑅𝑆𝑆/𝑛) where *k* is the number of parameters, *n* is the number of samples, and RSS is the residual sum of squares^27^.

### Geographic model

Sea surface temperatures were obtained from NASA’s earth observatory in an 8-day and 0.1 degrees resolution. Temperatures are given in 2,160 by 4,320 pixel resolution, where each pixel is 0.0833 degrees in latitude and 0.0833 degrees in longitude. For each marine metagenomic sample, the relevant pixel was found based on the sample coordinates. To obtain a robust measurement the temperature was estimated based on the median of the relevant pixel, and its 8 neighbors) and averaged on the measurements of the relevant 8-day window and the preceding 8-day window. Namely (s1+s2)/2, where s1 is the median of the 9 temperature values from the matching 8-day and s2 is the median of the 9 temperature values from the preceding 8 days. We then built a ridge regression model for the inference of sample temperatures based on the derived surface temperatures, along with the sample’s depth and latitude. We included first, second and third degrees of the depth in order to account for higher complexity in the effect of depth on temperature, as well as the absolute value and second degree of latitude in order to account for distance from the equator. We also included all pairwise interactions between elements of the three categories (surface temperature, depth, latitude). Hence the final model is: 𝑇∼𝑠 + 𝑑 + 𝑑^2^ + 𝑑^3^ + |𝑙| + 𝑙^2^ + 𝑠𝑑 + 𝑠𝑑^2^ + 𝑠𝑑^3^ + 𝑠|𝑙| + 𝑠𝑙^2^ + 𝑑|𝑙| + 𝑑𝑙^2^ + 𝑑^2^|𝑙| + 𝑑^2^𝑙^2^ + 𝑑^3^|𝑙| + 𝑑^3^𝑙^2^, where 𝑇 is the sampling temperature, 𝑠 is the matched surface temperature, 𝑑 is the sample’s depth, and 𝑙 is the sample’s latitude. We trained the model using LOSO cross-validation and compared the inferred temperatures to the samples’ real temperature measurements (**Figure S4**).

### Phylogeny association test

We used gene annotations from the OM-RGC9 and GMGC4 gene datasets to assign each read aligned to these datasets to a phylum or class. Taxa inclusion criteria required at least 30 valid samples per dataset, with a sample considered valid if it contained ≥5,000 reads assigned to the taxon. 4-mer frequencies were calculated using the first 5,000 reads per sample-taxon pair. Temperature inference was performed using Bayesian Ridge regression for each taxon. For comparison, we created a null distribution by randomly sampling 5,000 reads from each sample, following the same dataset distribution (i.e., the same number of samples from each of the metagenomic datasets), and repeated this process 1,000 times. This method controlled for sample size, dataset composition, and uneven representation of soil and marine environments.

### Codon analysis

For every mapped read in each sample, we counted codons according to the catalog-gene it was aligned to. Reads with indels or with more than 7 mismatches (5% of read length) were discarded. Any codons with degenerate nucleotide annotations (e.g., Ns representing any nucleotide or Y and R assigned to pyrimidine and purine) were also dismissed. The frequency of a codon within a sample was then calculated as the sum of counts of that codon across all reads, divided by the total codon counts in the sample.

#### Codon GC content correlation analysis

For each amino acid, we calculated the ratio between the frequency of the codons with the highest GC content and the sum of frequencies of all codons encoding that amino acid (GC codon bias). Spearman correlations were then calculated between these fractions and sample temperatures. After Bonferroni correction, the significance cut-off was p < 2.5×10^−4^.

To determine the confounding effect of nutrient availability, we performed three additional analyses:

1. We performed a two-step residual analysis by fitting a linear regression of temperature against nitrate, phosphate, or depth, and calculating the residuals, representing the variation in temperature not explained by the environmental covariate. These residuals were then centered by adding the mean temperature to maintain the original scale and correlated with the GC codon bias.
2. An Analysis of Covariance (ANCOVA) using Type III sums of squares was conducted. Type III sums of squares was chosen as it tests each effect after controlling for all other effects, providing a conservative estimate of significance that is not affected by unbalanced data and is independent of the order of effects in the model. The analysis was performed using Python’s statsmodels package (version 0.13.5), fitting a linear model with temperature as the dependent variable, the feature of interest as the independent variable, and the environmental covariate (nitrate, phosphate or depth) as the continuous covariate. The significance of each effect was evaluated using F-tests based on Type III sums of squares, with statistical significance set at p < 0.05 (**Table S3**).
3. We performed a focused analysis using only samples within nutrient ranges for which temperature was not correlated with the nutrient. We searched for nutrient ranges that (a) had at least 50 points; (b) encompassed a nutrient concentration range smaller than 1% of the entire environmental range in these samples (e.g. if nitrate encompassed a range of 0-40 µmol/L, the maximal range we considered was 0.4 µmol/L); and (c) temperature was uncorrelated with the nutrient (p-value>0.5 using Spearman’s rho p-value). The ranges are depicted in **Table S2**. In each of these ranges we calculated the GC codon bias correlation with temperature.

## Author contributions

T.A., O.L.-E. and D.Z. designed experiments, analyzed results and wrote the manuscript. T.Y. and Y.B. designed experiments, analyzed results and reviewed the manuscript. O.L.-E. and D.Z. conceived and supervised the project.

## Acknowledgements

We thank the Zeevi lab, Tal Korem, Shaul Pollak and Dana Bar-Zvi for fruitful discussions. This study was partly supported by the de Botton Center for Marine Science. O.L.-E. Is supported by the Rothschild postdoctoral fellowship and by EMBO postdoctoral fellowship. D.Z. is supported by the Alon Fellowship for the Integration of Outstanding Faculty.

## Supplementary Information

**Table S1: Total curated metadata and measurements of environmental parameters**

**Table S2:**
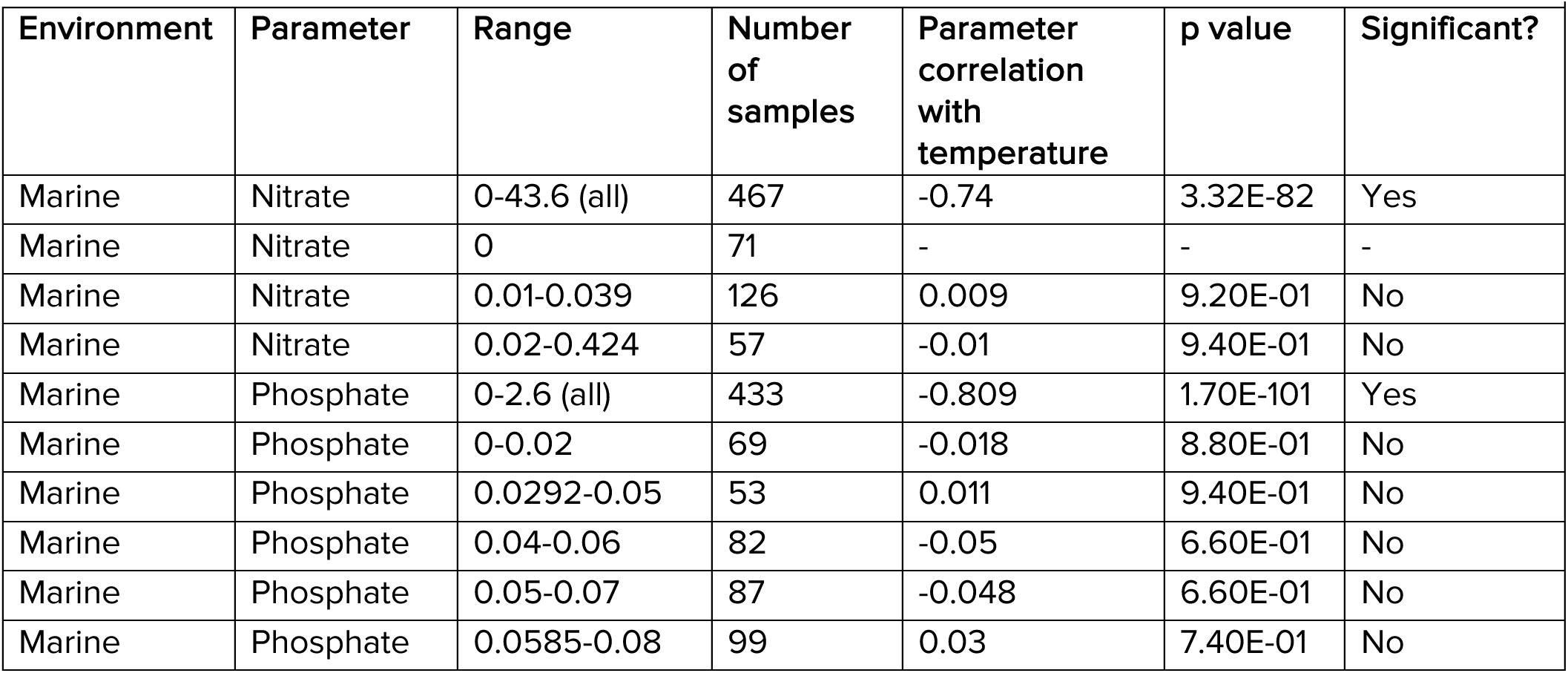
Correlations between temperature and different parameters in selected ranges.

| Environment | Parameter | Range | Number of samples | Parameter correlation with temperature | p value | Significant? |
| --- | --- | --- | --- | --- | --- | --- |
| Marine | Nitrate | 0-43.6 (all) | 467 | -0.74 | 3.32E-82 | Yes |
| Marine | Nitrate | 0 | 71 | - | - | - |
| Marine | Nitrate | 0.01-0.039 | 126 | 0.009 | 9.20E-01 | No |
| Marine | Nitrate | 0.02-0.424 | 57 | -0.01 | 9.40E-01 | No |
| Marine | Phosphate | 0-2.6 (all) | 433 | -0.809 | 1.70E-101 | Yes |
| Marine | Phosphate | 0-0.02 | 69 | -0.018 | 8.80E-01 | No |
| Marine | Phosphate | 0.0292-0.05 | 53 | 0.011 | 9.40E-01 | No |
| Marine | Phosphate | 0.04-0.06 | 82 | -0.05 | 6.60E-01 | No |
| Marine | Phosphate | 0.05-0.07 | 87 | -0.048 | 6.60E-01 | No |
| Marine | Phosphate | 0.0585-0.08 | 99 | 0.03 | 7.40E-01 | No |

**Table S3:** Summary of ANCOVA using type III sums of squares for the effect of environmental parameters on amino acid GC fraction.

| Feature | Depth (n=597) |  |  |  | Nitrate (n=467) |  |  |  | Phosphate (n=433) |  |  |  |
| --- | --- | --- | --- | --- | --- | --- | --- | --- | --- | --- | --- | --- |
|  | Feature F | Feature PR(>F) | Depth F | Depth PR(>F) | Feature F | Feature PR(>F) | Nitrate F | Nitrate PR(>F) | Feature F | Feature PR(>F) | Phosphate F | Phosphate PR(>F) |
| A | 160.77 | 8.94E-33 | 5.23 | 2.26E-02 | 8.16 | 4.47E-03 | 165.61 | 1.27E-32 | 1.27 | 2.60E-01 | 454.18 | 2.62E-69 |
| C | 96.01 | 4.10E-21 | 11.28 | 8.35E-04 | 2.62 | 1.06E-01 | 219.21 | 6.75E-41 | 4.09 | 4.37E-02 | 485.32 | 1.51E-72 |
| D | 99.18 | 1.04E-21 | 5.72 | 1.71E-02 | 0.76 | 3.83E-01 | 227.08 | 4.68E-42 | 1.60 | 2.07E-01 | 538.46 | 7.94E-78 |
| E | 16.66 | 5.10E-05 | 30.93 | 4.04E-08 | 2.73 | 9.89E-02 | 368.61 | 6.83E-61 | 0.01 | 9.37E-01 | 605.52 | 4.34E-84 |
| F | 0.37 | 5.46E-01 | 30.07 | 6.17E-08 | 19.82 | 1.07E-05 | 400.09 | 1.22E-64 | 5.12 | 2.42E-02 | 626.98 | 5.24E-86 |
| G | 229.91 | 3.87E-44 | 2.71 | 1.00E-01 | 21.44 | 4.75E-06 | 141.51 | 1.15E-28 | 4.36 | 3.73E-02 | 406.25 | 4.31E-64 |
| H | 107.82 | 2.53E-23 | 10.61 | 1.19E-03 | 1.97 | 1.61E-01 | 226.51 | 5.66E-42 | 0.65 | 4.19E-01 | 546.76 | 1.26E-78 |
| I | 41.46 | 2.48E-10 | 7.49 | 6.39E-03 | 2.28 | 1.32E-01 | 272.54 | 1.68E-48 | 1.87 | 1.73E-01 | 526.79 | 1.08E-76 |
| K | 24.98 | 7.64E-07 | 19.65 | 1.11E-05 | 0.10 | 7.58E-01 | 334.31 | 1.22E-56 | 0.05 | 8.16E-01 | 576.64 | 1.92E-81 |
| L | 54.28 | 5.85E-13 | 5.07 | 2.47E-02 | 1.93 | 1.65E-01 | 251.53 | 1.42E-45 | 0.66 | 4.16E-01 | 498.61 | 6.77E-74 |
| N | 94.44 | 8.11E-21 | 7.43 | 6.62E-03 | 1.29 | 2.57E-01 | 239.39 | 7.64E-44 | 3.68 | 5.58E-02 | 564.78 | 2.46E-80 |
| P | 116.69 | 5.89E-25 | 2.30 | 1.30E-01 | 0.98 | 3.23E-01 | 188.90 | 2.65E-36 | 0.01 | 9.40E-01 | 461.54 | 4.39E-70 |
| Q | 42.67 | 1.39E-10 | 4.65 | 3.14E-02 | 2.97 | 8.56E-02 | 275.85 | 5.91E-49 | 2.96 | 8.59E-02 | 557.10 | 1.31E-79 |
| R | 182.33 | 1.99E-36 | 1.62 | 2.04E-01 | 11.99 | 5.85E-04 | 150.00 | 4.45E-30 | 1.60 | 2.07E-01 | 424.49 | 4.11E-66 |
| S | 107.12 | 3.41E-23 | 3.58 | 5.91E-02 | 0.94 | 3.32E-01 | 205.15 | 8.58E-39 | 0.32 | 5.74E-01 | 470.73 | 4.82E-71 |
| T | 172.91 | 7.62E-35 | 3.04 | 8.16E-02 | 9.10 | 2.69E-03 | 160.90 | 7.31E-32 | 1.60 | 2.06E-01 | 441.41 | 6.02E-68 |
| V | 112.43 | 3.55E-24 | 4.15 | 4.22E-02 | 1.82 | 1.78E-01 | 199.17 | 6.97E-38 | 0.13 | 7.15E-01 | 474.44 | 1.99E-71 |
| Y | 63.95 | 6.68E-15 | 10.85 | 1.05E-03 | 0.00 | 9.99E-01 | 261.87 | 5.03E-47 | 1.24 | 2.66E-01 | 558.20 | 1.03E-79 |

**Figure S1:**
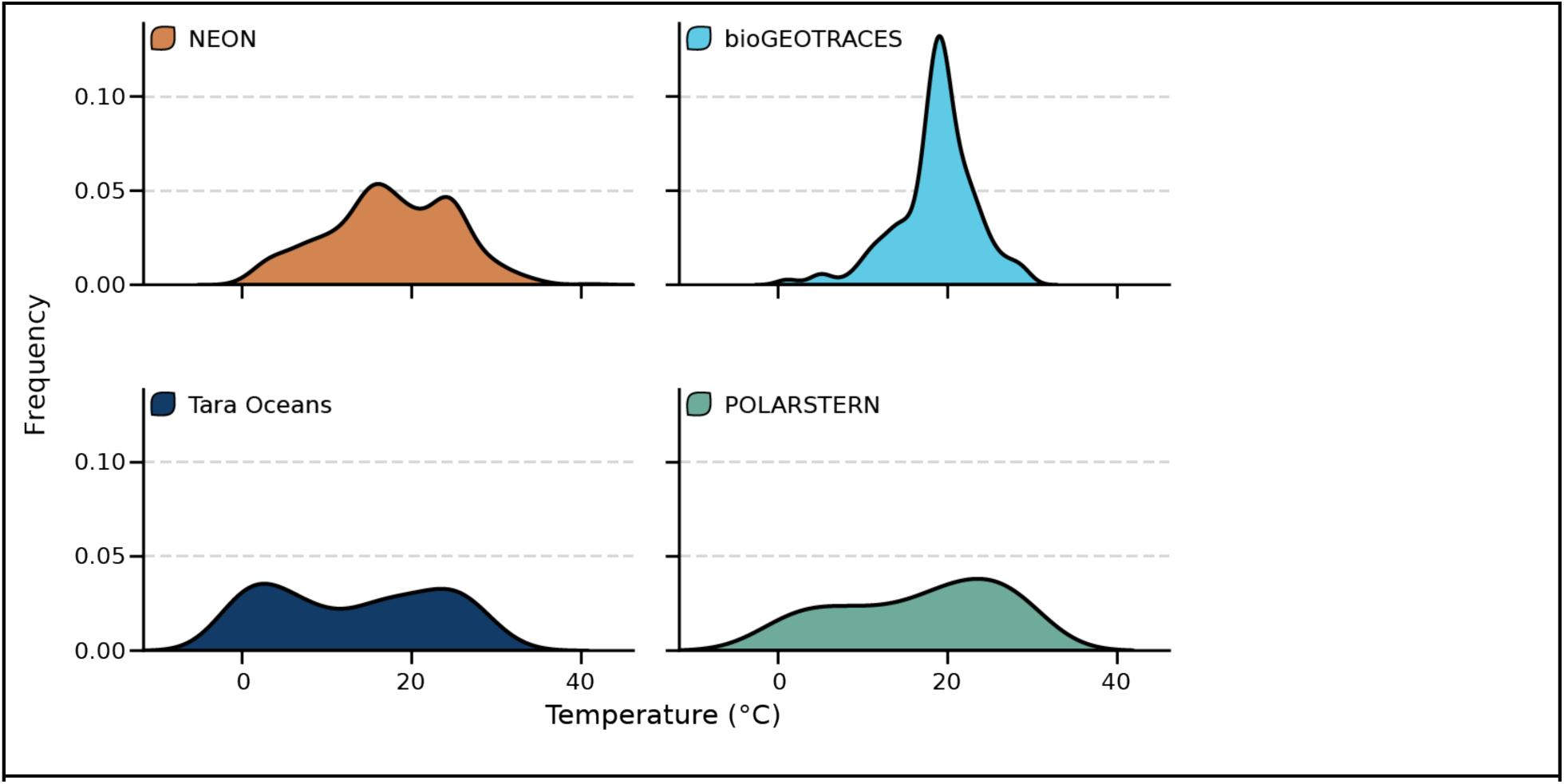
Frequency distributions of sample temperatures, grouped and colored by dataset.

**Figure S2:**
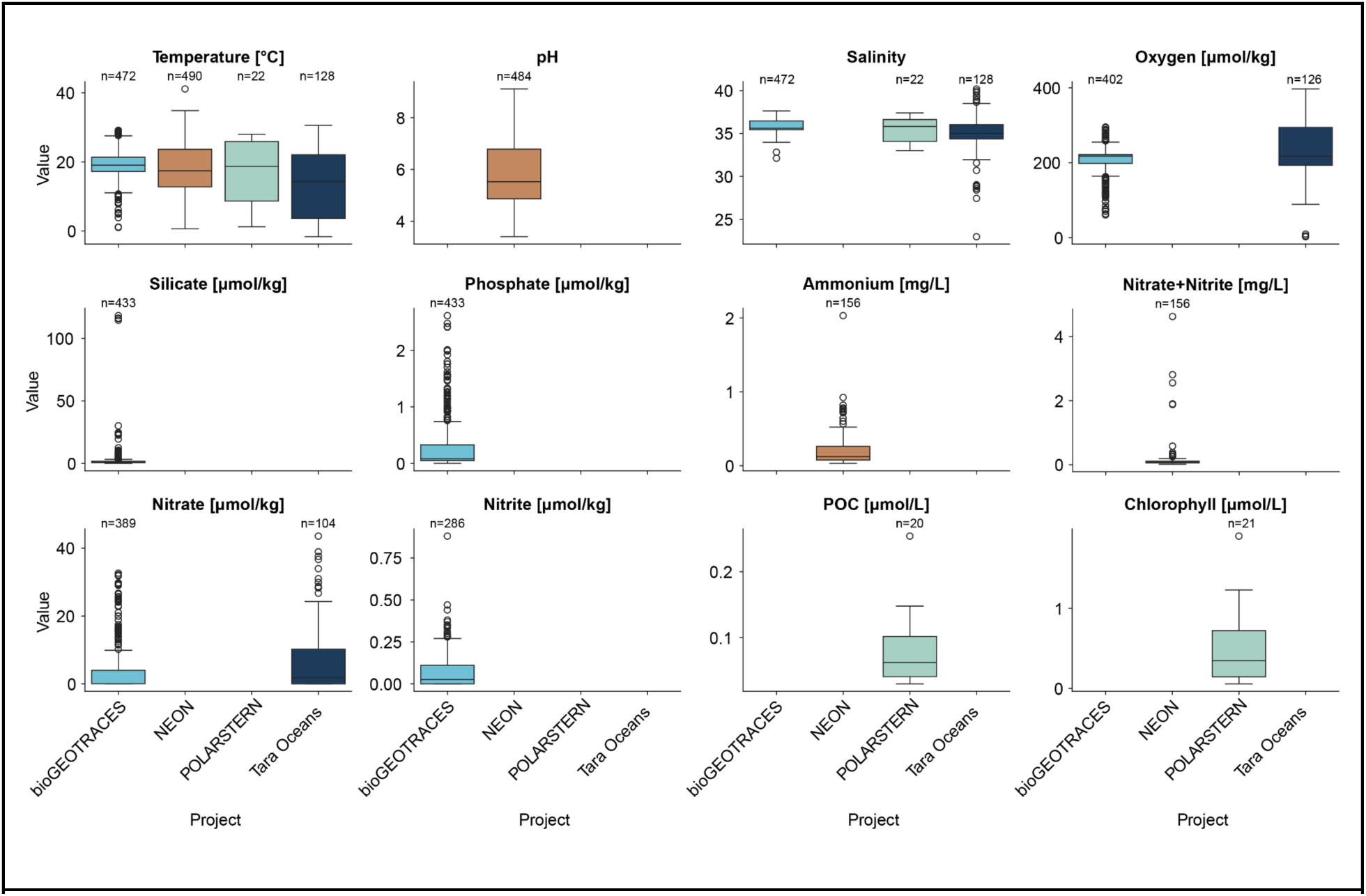
Distributions of other parameters that appeared in the datasets.

**Figure S3:**
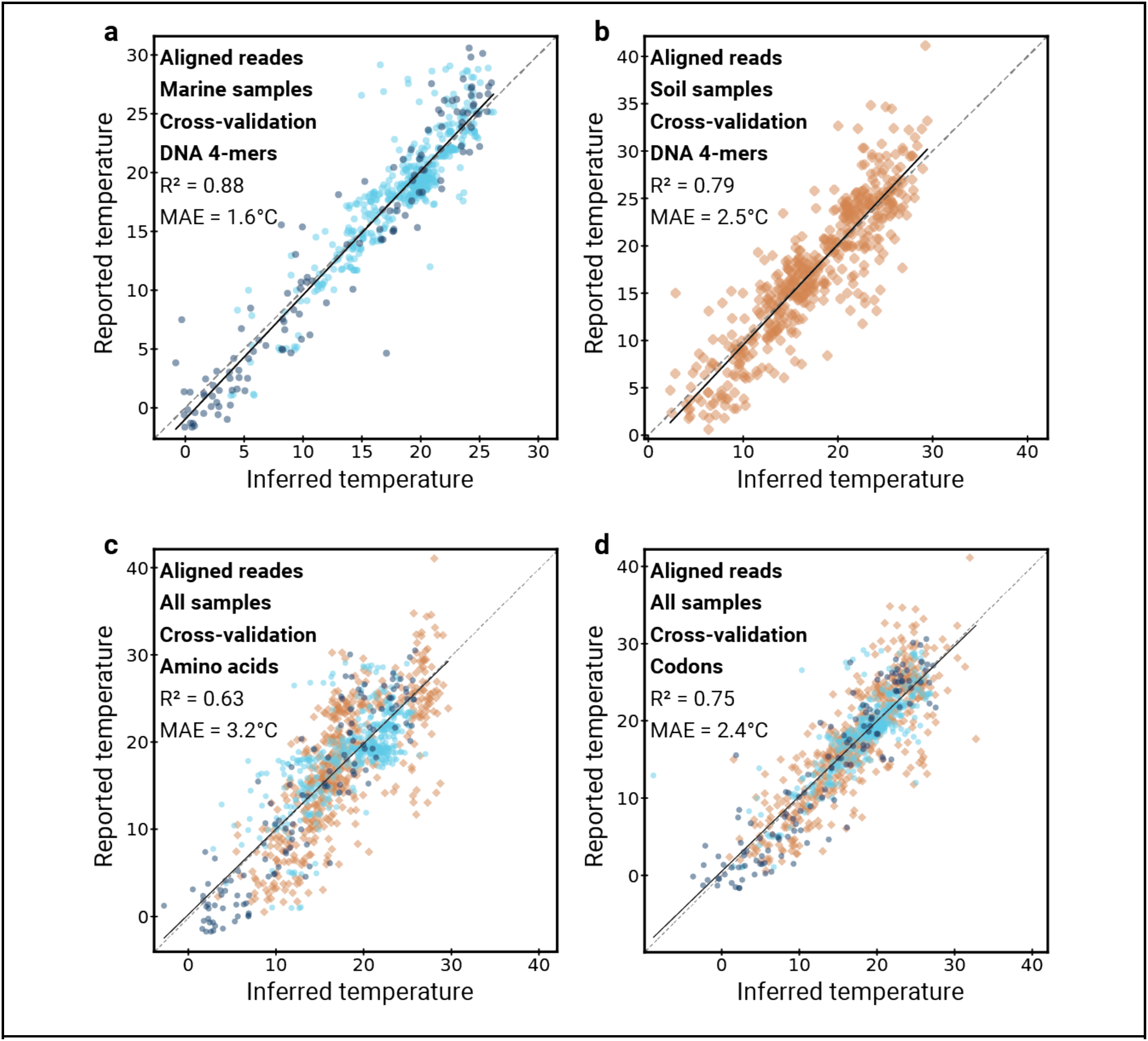
Inference comparison (Ridge regression, Cross-validation, 1M aligned reads): **[a]** DNA 4-mers, marine samples. **[b]** DNA 4-mers, soil samples. **[c]** Amino acid frequencies, all samples. **[d]** Codon frequencies, all samples.

**Figure S4:**
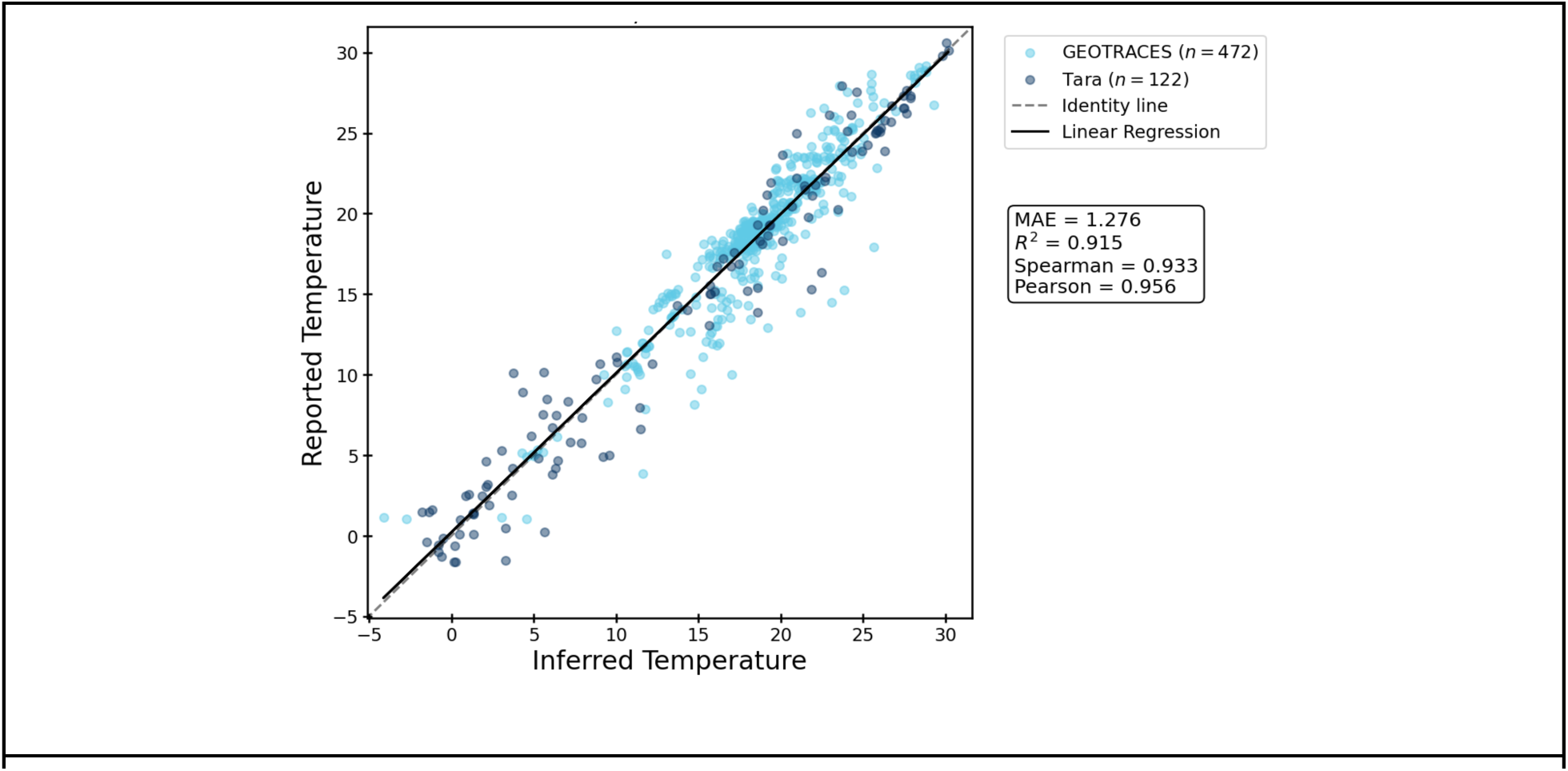
Temperature inference based on a model that relies on latitude, depth, and temperature at the surface level (collected from NASA). Using the formula: 𝑇∼𝑠 + 𝑑 + 𝑑^2^ + 𝑑^3^ + |𝑙| + 𝑙^2^ + 𝑠𝑑 + 𝑠𝑑!^2^+ 𝑠𝑑^3^ + 𝑠|𝑙| + 𝑠𝑙^2^ + 𝑑|𝑙| + 𝑑𝑙^2^ + 𝑑^2^|𝑙| + 𝑑^2^𝑙^2^ + 𝑑^3^|𝑙| + 𝑑^3^𝑙^2^. 𝑇 is the sampling temperature, 𝑠 is the matched surface temperature, 𝑑 is the sample’s depth, and 𝑙 is the sample’s latitude.

**Figure S5:**
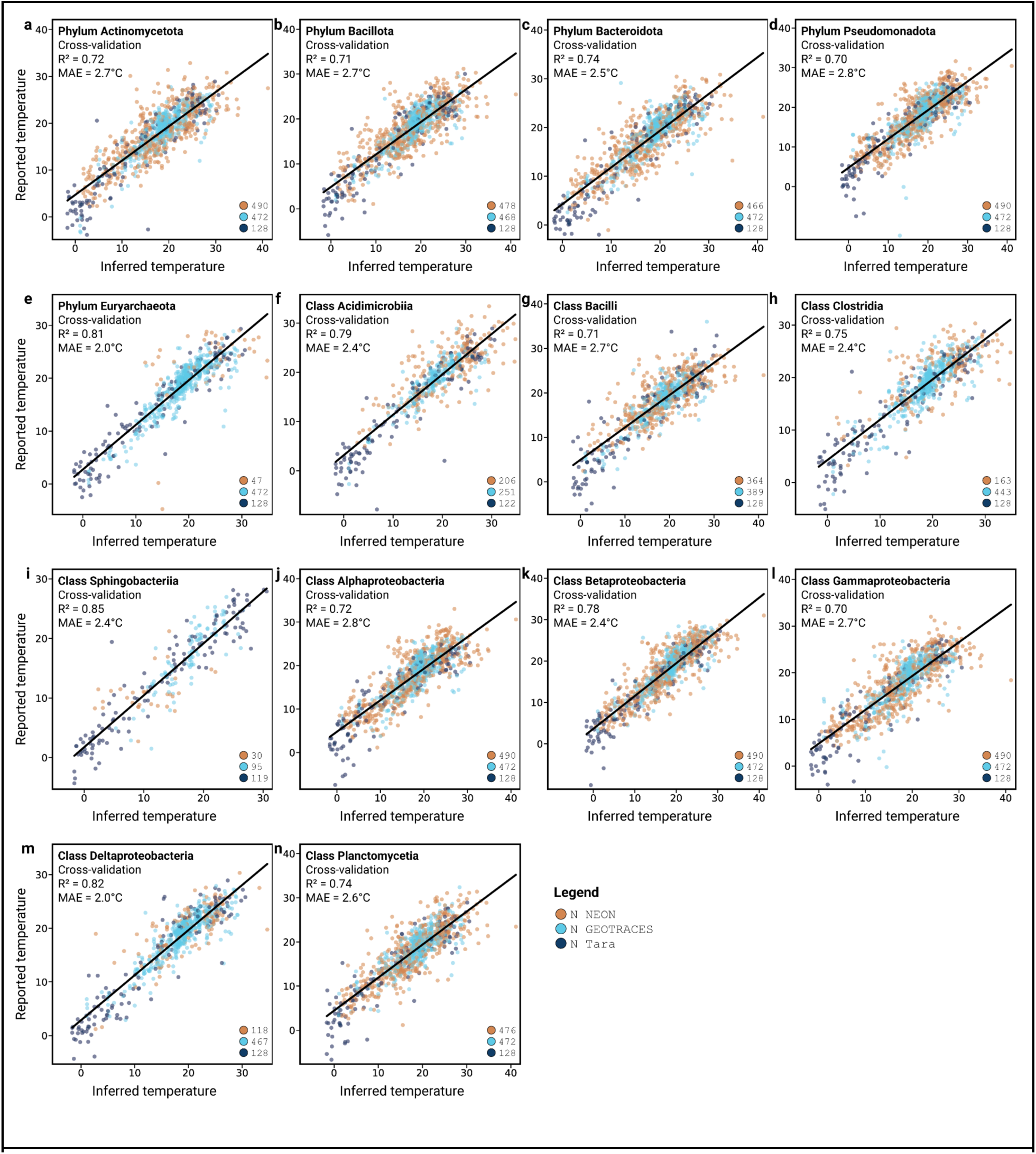
**[a-n]** Scatter plots of inferred versus reported temperatures using cross-validation and 4-mers from metagenomic reads assigned to taxa presented in **Fig. 2**.

**Figure S6:**
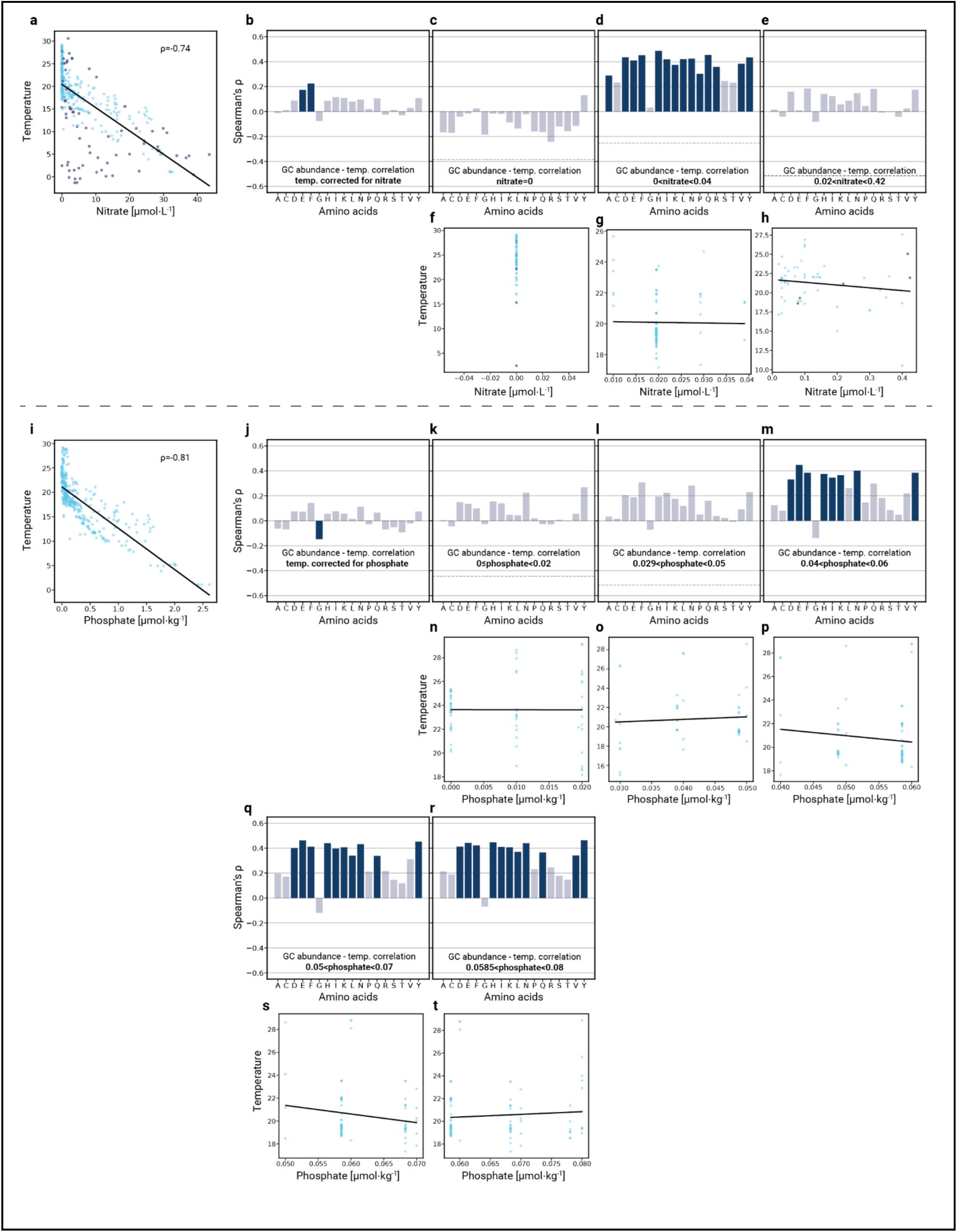
Correlations between GC-rich codons and temperature alter in specific ranges. **[a]** and **[i]** Temperatures in marine samples (GEOTRACES in light blue and Tara in dark blue) plotted against nitrate concentrations and phosphate concentrations respectively. **[b-e]** and **[j-m, q-r]** Spearman correlations between temperature and GC codon bias. **[f-h]** Scatter plots of temperature versus nitrate concentrations in specific nitrate ranges, matching the range in the bar plot above them. **[n-p, s-t]** Same as **[f-h]** but for phosphate. **[c-e]** Correlations calculated in specific nitrate ranges. **[k-m, q-r]** Same as **[c-e]** but for phosphate. Nitrate ranges (μmol/L; non-inclusive unless stated): **[c,f]** range: 0 (inclusive, n=71); **[d,g]** range: 0-0.04 (n=126); **[e,h]** range: 0.02-0.42 (n=57). Phosphate ranges (μmol/kg; inclusive unless stated): **[k,n]** range: 0-0.02 (n=69); **[l,o]** range: 0.029-0.05 (n=53); **[m,p]** range: 0.04-0.06 (n=82); **[q,s]** range: 0.05-0.07 (n=87); **[r,t]** range: 0.0585-0.08 (n=99).

